# The evolution of parental effects when selection acts on fecundity versus viability

**DOI:** 10.1101/741835

**Authors:** Bram Kuijper, Rufus A. Johnstone

**Affiliations:** Centre for Ecology & Conservation, College of Life and Environmental Sciences, University of Exeter, Penryn TR10 9FE, United Kingdom; Behaviour and Evolution Group, Department of Zoology, University of Cambridge, Downing Street, Cambridge CB2 3EJ, United Kingdom

**Keywords:** epigenetics, maternal effects, nongenetic inheritance, phenotypic plasticity, information, transgenerational effects, environmental stochasticity

## Abstract

Most predictions on the evolution of adaptive parental effects and phenotypic memory exclusively focus on the role of the abiotic environment. How parental effects are affected by population demography and life history is less well understood. To overcome this, we use an analytical model to assess whether selection acting on fecundity versus viability affects the evolution of parental effects in a viscous population experiencing a spatiotemporally varying environment. We find that parental effects commonly evolve in regimes of viability selection, but are less likely to evolve in regimes of fecundity selection. In regimes of viability selection, an individual’s phenotype becomes correlated with its local environment during its lifetime, as those individuals with a locally adapted phenotype are more likely to survive until parenthood. Hence, a parental phenotype rapidly becomes an informative cue about its local environment, favoring the evolution of parental effects. By contrast, in regimes of fecundity selection, locally maladapted and adapted parents survive at equal rates, so that the parental phenotype, by itself, is not informative about the local environment. Correlations between phenotype and environment still arise, but only when more fecund, locally adapted individuals leave more successfully established offspring to the local patch. Hence, correlations take at least two generations to develop, making them more sensitive to distortion by environmental change or competition with immigrant offspring. Hence, we conclude that viability selection is most conducive to the evolution of adaptive parental effects in spatially structured populations.

## 1 Introduction

An accumulating number of studies shows that parents provide a range of inputs that contribute to offspring development (Uller, 2008; Jablonka & Raz, 2009; Danchin *et al.*, 2011; Heard & Martienssen, 2014; Chen *et al.*, 2016). These parental effects can be mediated by a variety of mechanisms, such as maternal hormones (Meylan *et al.*, 2012), the transmission of chromatin modifications and epimutations (Heard & Martienssen, 2014), maternal nutrients (English & Uller, 2016) or the transmission of behavior via social learning (Mesoudi *et al.*, 2016). Crucially, parental effects result in transgenerational plasticity, in which the phenotype of an offspring is not only dependent on the genes it inherits and its current environment, but also dependent on the phenotypes or environments of its parents and grandparents (Uller, 2008; Herman & Sultan, 2011; Burton & Metcalfe, 2014). In order to understand when and where transgenerational plasticity is likely to be important, predictions are needed about the ecological and social conditions that favor the evolution of parental effects.

Indeed, a series of novel models have started to consider the evolution of parental effects, finding that they are particularly likely to evolve in fluctuating environments in which parental and offspring environments are correlated (e.g., Rivoire & Leibler, 2014; Kuijper *et al.*, 2014; Uller *et al.*, 2015; McNamara *et al.*, 2016; Proulx & Teotónio, 2017) and when this environment imposes strong selection (English *et al.*, 2015; Kuijper & Hoyle, 2015). While these models provide the field with novel and testable predictions (e.g., see Dey *et al.*, 2016), a shortcoming is their exclusive focus on the abiotic environment. By contrast, it is currently poorly understood whether other, biotic, factors such as demography and life history are also of importance when predicting the evolution of transgenerational plasticity (Kuijper & Johnstone, 2016).

One major point of focus in life history theory are the differential consequences of fecundity versus viability selection. In certain cases, adapted individuals produce larger numbers of offspring than other individuals (fecundity selection; Pincheira-Donoso & Hunt, 2017), whereas in other cases such individuals instead experience higher levels of survival (viability selection). Here we therefore ask whether the evolution of parental effects is differentially affected by fecundity versus viability selection.

Previous work on the evolution of mutation rates (Leigh, 1973; Ishii *et al.*, 1989) and stochastic phenotype switching (Acar *et al.*, 2008; Salathé *et al.*, 2009; Gaál *et al.*, 2010; Carja *et al.*, 2014) in fluctuating environments gives us little indication that the mode of selection matters at all to the evolution of inheritance mechanisms: selection in these models can represent selection on fecundity or viability without much difference. Only when selection acts on combinations of male and female genotypes (fertility selection in the classical population genetic sense: Travis, 1988), have effects on the mutation rate been found (Holsinger *et al.*, 1986). However, when selection results in individual differences in fecundity regardless of partner phenotype, the mode of selection is irrelevant.

As these previous studies focus exclusively on well-mixed populations, it raises the question whether fecundity and viability selection still lead to equivalent results when relatives interact. A recent model on the evolution of parental effects in viscous populations shows that interactions among relatives are highly conducive to the evolution of parental effects (Kuijper & Johnstone, 2016): parental effects allow mothers to produce offspring with more heterogeneous phenotypes relative to mothers that exclusively rely on genetic inheritance. Such variability is highly advantageous at the level of the group, as it makes it more likely that at least some offspring survive environmental change (Moran, 1992; McNamara, 1995; Leimar, 2005; Lehmann & Balloux, 2007). However, as the model of Kuijper & Johnstone (2016) exclusively considered viability selection, it remains to be seen how robust these results are to the mode of selection. From studies that focus on the evolution of helping and altruism, we already know that fecundity and viability selection lead to differences in the structure of competition among kin (Pen, 2000; Taylor & Irwin, 2000; Lion & Gandon, 2010; Taylor, 2010), which could potentially also affect the evolution of parental effects.

To address the role of fecundity versus viability selection on the evolution of parental effects, we build on previous models (Kuijper & Johnstone, 2016; Leimar & McNamara, 2015; Leimar *et al.*, 2016; English & Uller, 2016) by considering phenotypic development as a ‘switching device’, which integrates cues from the offspring’s genes and the parental phenotype as inputs. For the sake of clarity, the current study focuses on the relative importance of genetic versus parental cues, and we refer the reader to other recent models that have considered environmental cues in the current or the parental generation (see Kuijper & Hoyle, 2015; McNamara *et al.*, 2016; English *et al.*, 2015; Proulx & Teotónio, 2017). Similar to previous models, we consider an individual’s phenotype as a binary variable, so the switching device decides on producing one or the other of two alternative morphs that each match a different environment (Roff, 1996; Leimar *et al.*, 2006; Chevin & Lande, 2013). Over evolutionary time, the characteristics of the switching device can be altered by the successive substitution of mutant modifier alleles (see also Salathé *et al.*, 2009; Carja *et al.*, 2014). Examples of such modifier alleles are alleles coding for a different offspring sensitivity to maternal hormones (Tschirren *et al.*, 2009), alter the regulatory specificity of certain small RNAs that are passed on through the gametes (Liebers *et al.*, 2014; Chen *et al.*, 2016) or change the epigenetic resetting rate (Heard & Martienssen, 2014; Taudt *et al.*, 2016).

## 2 The model

The model assumes a spatially structured population that is divided into infinitely many patches (Wright, 1931), which are linked by juvenile dispersal. Each patch has a fixed population size of *n* breeders and switches back and forth between two possible environmental states **e** = {*e*_1_, *e*_2_}. For reasons of tractability we restrict the analysis to *n* = 2 breeders per patch. Stochastic individual-based simulations show, however, that much larger numbers of breeders per patch (e.g., *n* = 25) result in qualitatively similar outcomes (see Supplementary Figure S4).

Individuals can adopt one of two possible phenotypes **z** = {*z*_1_, *z*_2_} making them locally adapted when their phenotype *z*_*i*_ matches the environmental state *e*_*i*_ of the local patch. Within this fluctuating environment, we then focus on the evolution of a genetically determined strategy (*p*_1_, *p*_2_), the elements of which determine the probability of producing a phenotype *z*_1_ offspring by *z*_1_ or *z*_2_ parent respectively. When these phenotype determination probabilities evolve to be independent of the parental phenotype (i.e., *p*_1_ = *p*_2_) no parental effects evolve and offspring phenotypes are considered to be genetically determined. However, when *p*_1_ ≠*p*_2_, parental effects arise as the offspring phenotype determination is a function of the parent’s phenotype. Note also that our model focuses on parental effects based on the parental phenotype, rather than the maternal environment or a mother’s genes. Hence this is an evolutionary model of a cascading maternal effect (McGlothlin & Galloway, 2013; Kuijper & Hoyle, 2015).

We employ an adaptive dynamics approach (Geritz *et al.*, 1998; McGill & Brown, 2007; Dercole & Rinaldi, 2008) to model the evolution of (*p*_1_, *p*_2_), assuming that evolutionary change occurs through the successive substitution of mutations that have small phenotypic effects. We assume a separation of the ecological and evolutionary timescales, implying that demographic changes (environmental change, deaths and births) occur at a much faster timescale than evolutionary change in the genetic modifiers that specify the probabilities of phenotypic inheritance. An analytical description of the model is provided in the Supplementary Mathematica Notebook.

### Population dynamics: environmental change and breeder viability/fecundity

The population dynamics of the model are described by a continuous-time dynamic, so that events occur one-at-a-time (Moran, 1958; Metz & Gyllenberg, 2001; Alizon & Taylor, 2008). The following two events can occur: i) environmental change, where a patch in environmental state *e*_*j*_ switches to environmental state *e*_*i*_ with rate 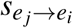. When the environment changes, all adapted phenotypes become maladapted and all maladapted individuals become adapted. 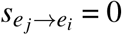 implies a constant environment, whereas *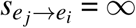* reflects an ever-changing environment. Importantly, *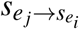* reflects the rate of change of an individual patch, rather than the global environment as a whole. Hence, at equilibrium, the total frequency of patches 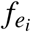 and 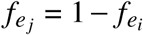 in state *e*_*i*_ and *e*_*j*_ is constant and given by 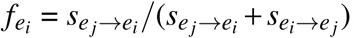. Because we vary both 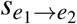 and *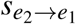* independently, we use the aggregate variable

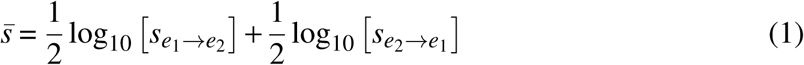

to reflect the global average rate of environmental change in the population (Kuijper & Johnstone, 2016).

ii) breeder viability: adult breeders with phenotype *z*_*i*_ in an *e* _*j*_ environment have a mortality rate given by 0 *< M*_*ij*_ *< ∞* where a breeder maladapted to its local environment has a higher mortality rate than an adapted breeder: *M*_*ii*_ *< M*_*ij*_. Prior to the occurrence of mortality event, all adult breeders produce offspring, with phenotype *z*_*i*_ breeders living in environment *e*_*j*_ produces having relative fecundity 0 *< B*_*ij*_ *< ∞*. The now vacant breeding position in an environment *e*_*j*_ is subsequently filled (maintaining a local population size of *n* = 2) with a juvenile offspring, as a result of competition among all locally born offspring remaining on their natal site and any immigrant offspring. Specifically, the probability that a newborn offspring born from a local adult breeder takes over the vacant patch is then given by 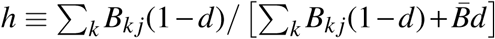. Here, *B*_*kj*_ is the fecundity of the *k*th adult breeder in the local *e*_*j*_ patch and 1 - *d* the probability that offspring remain at the natal patch. In addition, *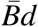* reflects the total number of immigrant offspring, where *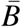* is the average fecundity in the global environment and *d* the juvenile dispersal probability.

### Evolutionary dynamics

We model evolution in the two phenotype determination strategies *p*_1_, *p*_2_, by considering the successive invasion of rare mutants that slightly differ in one of these three traits from the otherwise monomorphic resident population. In the Supplementary Mathematica notebook, we derive an expression of the instantaneous change in fitness 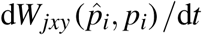 of an adult breeder who has phenotype *x ∈* {*z*_1_, *z*_2_}, lives in environment *j* and has a neighbour of phenotype *y ∈* {*z*_1_, *z*_2_}, while expressing the mutant strategy 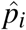 in a resident population with strategy *p*_*i*_. As shown in the Supplementary Mathematica Notebook, each of the events (deaths, births, environmental change) that affect the adult breeder are weighed by the respective gains and losses in terms of reproductive value of the mutant allele (e.g., Taylor, 1990, 1996; Härdling *et al.*, 2003; Alizon & Taylor, 2008; Wild *et al.*, 2009).

If evolution proceeds slowly, so that an individual’s lifespan represents only an infinitesimal fraction of evolutionary time, a standard result in adaptive dynamics (e.g., Dieckmann & Law, 1996) allows one to describe changes in character values over time using the following evolutionary dynamic (e.g., see eq. C5 in Härdling *et al.*, 2003)

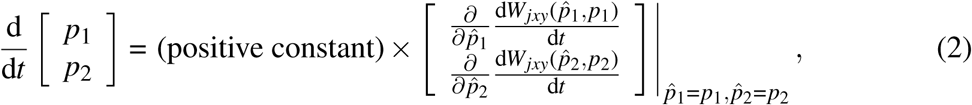

where we assume that any pleiotropic mutations are absent. We then find evolutionary endpoints by iterating the adaptive dynamic in (2) starting from a particular set of values [*p*_1_, *p*_2_]_*t*=0_ until it vanishes,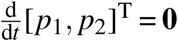. During each time step of the iteration, we numerically solved for equilibrium values of patch type frequencies, reproductive values and relatedness for the current values of [*p*_1_, *p*_2_]_*t*_ using a root finding algorithm written in C++. Convergence was determined when the largest difference in values of [*p*_1_, *p*_2_] between consecutive time steps was *≤*10^-7^. Starting values used in our iterations are [*p*_1_, *p*_2_]_*t*=0_ = [0.5, 0.5]. The outcomes obtained from these numerical iterations are convergence stable by definition, and individual-based simulations revealed that values are also evolutionarily stable. The Mathematica notebook as well as code for the numerical iterations and individual-based simulations is available at https://doi.org/10.5281/zenodo.3372450.

## 3 Results

We study parental effects by evolving the strategy vector (*p*_1_, *p*_2_), which specifies the proportion of *z*_1_-offspring produced by a parent with phenotype *z*_1_ or *z*_2_ respectively. If *p*_1_ = *p*_2_, offspring phenotypes are determined independently of a parent’s phenotype, so that parental effects are necessarily absent. When *p*_1_ ≠ *p*_2_ however, the offspring phenotype is a function of the parent’s phenotype. Figure 1 illustrates how fecundity and viability selection impact on the prevalence of parental effects, for an example case where juvenile dispersal is limited (*d* = 0.1) and the rate of environmental change is slow (*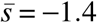*, see eq. [1]). Similar to previous analyses (Kuijper & Johnstone, 2016), we find that there are substantial regions where parental effects do not evolve at all (i.e., *p*_1_ = *p*_2_, white areas in Figure 1), particularly when one environment is much more common than the other. Here, parents of both phenotypes only produce the phenotype that matches the most prevalent environment.

**Figure 1:**
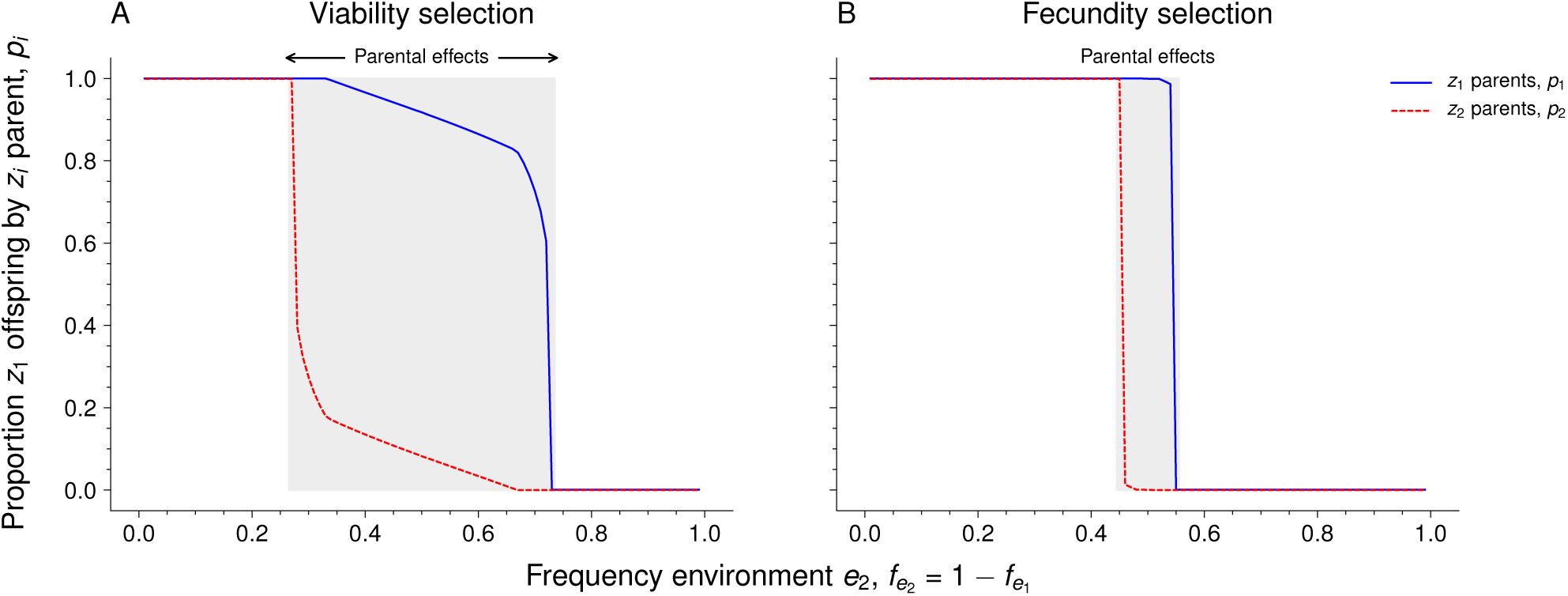
Parental effects (grey areas where *p*_1_ ≠ *p*_2_) are much more likely to occur in regimes of viability selection (panel A) than fecundity selection (panel B). Parameters: *M*_*a*_ = 1, *M*_*m*_ = 2, *B*_*a*_ = *B*_*m*_ = 1 (panel A), *M*_*a*_ = *M*_*m*_ = 1, *B*_*a*_ = 2, *B*_*m*_ = 1 (panel B), *d* = 0.1, 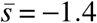.

By contrast, when the frequencies of the two environments are roughly similar (i.e., around the point 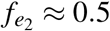), we find that *p*_1_ ≠ *p*_2_, so that offspring phenotype determination now depends on the parental phenotype. Interestingly, the scope for parental effects appears to be much larger in regimes of viability selection relative to regimes of fecundity selection. Moreover, this difference between viability selection and fecundity selection appears to be relatively robust to larger number of breeders per patch *n*, where parental effects evolve in contexts of viability selection (but not in contexts of fecundity selection) up to *n* = 50 (Figure S4).

### General ecological conditions favoring parental effects

Next, we assess whether this differential effect of fecundity and viability selection is robust to changes in juvenile dispersal *d* or the average rate of environmental change 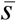. Figure 2 depicts the expected number of consecutive generations over which phenotype *z*_1_ is inherited from parent to offspring, while (i) varying the frequencies of the two environments, (ii) the average rate of environmental change and (iii) the rate of dispersal (the duration of inheritance of phenotype *z*_2_ is a mirror image of Figure 2, see Figure S2).

**Figure 2:**
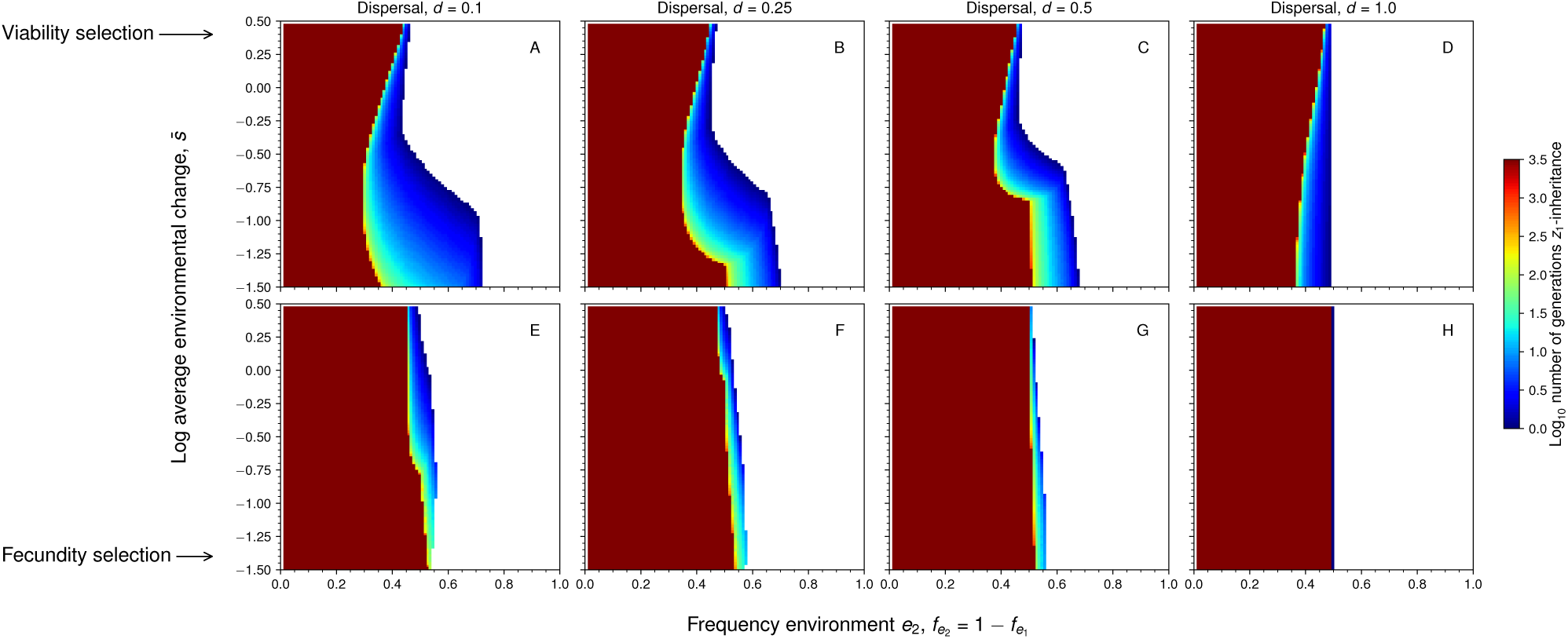
Viability selection (panels A-D) is much more likely to lead to intermediate fidelities of inheritance (characteristic of parental effects) of phenotype *z*_1_ (orange to blue colored areas) than fecundity selection (panels E-H). Dark-red areas indicate high-fidelity inheritance of phenotype *z*_1_, which occurs in populations monomorphic for *z*_1_, implying that parental effects are absent. Parameters: *M*_*a*_ = 1, *M*_*m*_ = 2, *B*_*a*_ = *B*_*m*_ = 1 (panels A-D), *M*_*a*_ = *M*_*m*_ = 1, *B*_*a*_ = 2, *B*_*m*_ = 1 (panels E-H), *n* = 2. See Figure S1 for the inheritance of phenotype *z*_2_, which is a mirror image of the current figure.

Similar to Figure 1, Figure 2 shows that when one environment predominates (dark red regions at the left or right hand sides of each panel), no parental effects evolve regardless of the presence of fecundity or viability selection. Rather, populations are monomorphic for the phenotype favored in the most prevalent environment (*z*_1_ when *e*_2_ is rare, *z*_2_ when *e*_2_ is common), resulting in a monomorphism and long-term (genetic) inheritance.

Only when both environments are encountered at roughly similar rates (middle of each panel in Figure 2), do we find that the parental effects evolve, as the inheritance spans only a limited number of generations. However, comparing Figures 2A-D with Figures 2E-H shows that parental effects are most commonly encountered in populations experiencing viability selection. By contrast, in populations which experience fecundity selection parental effects only evolve when populations encounter both environments at almost exactly the same rate.

Next, Figure 2 shows that parental effects are more likely to evolve in populations with limited dispersal, corroborating results from previous studies (e.g., English *et al.*, 2015; Leimar & McNamara, 2015; Kuijper & Johnstone, 2016), albeit with two exceptions. First, we find that the positive effect of limited dispersal on parental effects is much weaker in populations experiencing fecundity selection. Second, in populations experiencing viability selection, we find a limited number of cases where parental effects even evolve when all juveniles disperse to (random) remote patches (*d* = 1, Figure 2D). To understand this surprising result, Supplementary Figure S2 shows that viability selection favors a global mixture of *z*_1_ and *z*_2_ offspring, which can be achieved through many alternative combinations of *p*_1_ and *p*_2_ (i.e., a line of equilibria), where typically *p*_1_ ≠ *p*_2_ thus resulting in parental effects. By contrast, in populations with fecundity selection and *d* = 1, a monomorphism of the phenotype that matches the commonest environment is always selectively favored, selecting against any parental effects.

### Interactions between fecundity and viability selection

Finally, Figure 3 explores the evolution of parental effects when populations experience fecundity and viability selection simultaneously. In populations where viability selection prevails, increasing the strength of fecundity selection has little effect (Figure 3A). If anything, increasing levels of fecundity selection seem to enhance rather than inhibit the evolution of parental effects, at least when viability selection is already present. In populations where fecundity selection prevails (Figure 3B), increasing the effect of viability selection substantially increases the scope for parental effects. Hence, we conclude that fecundity selection does not reduce the scope for parental effects in populations which also experience viability selection; only when viability selection is weak, is fecundity selection most likely to limit the evolution of parental effects. A more general overview of the interaction between fecundity and viability selection is given in Figure S3.

**Figure 3:**
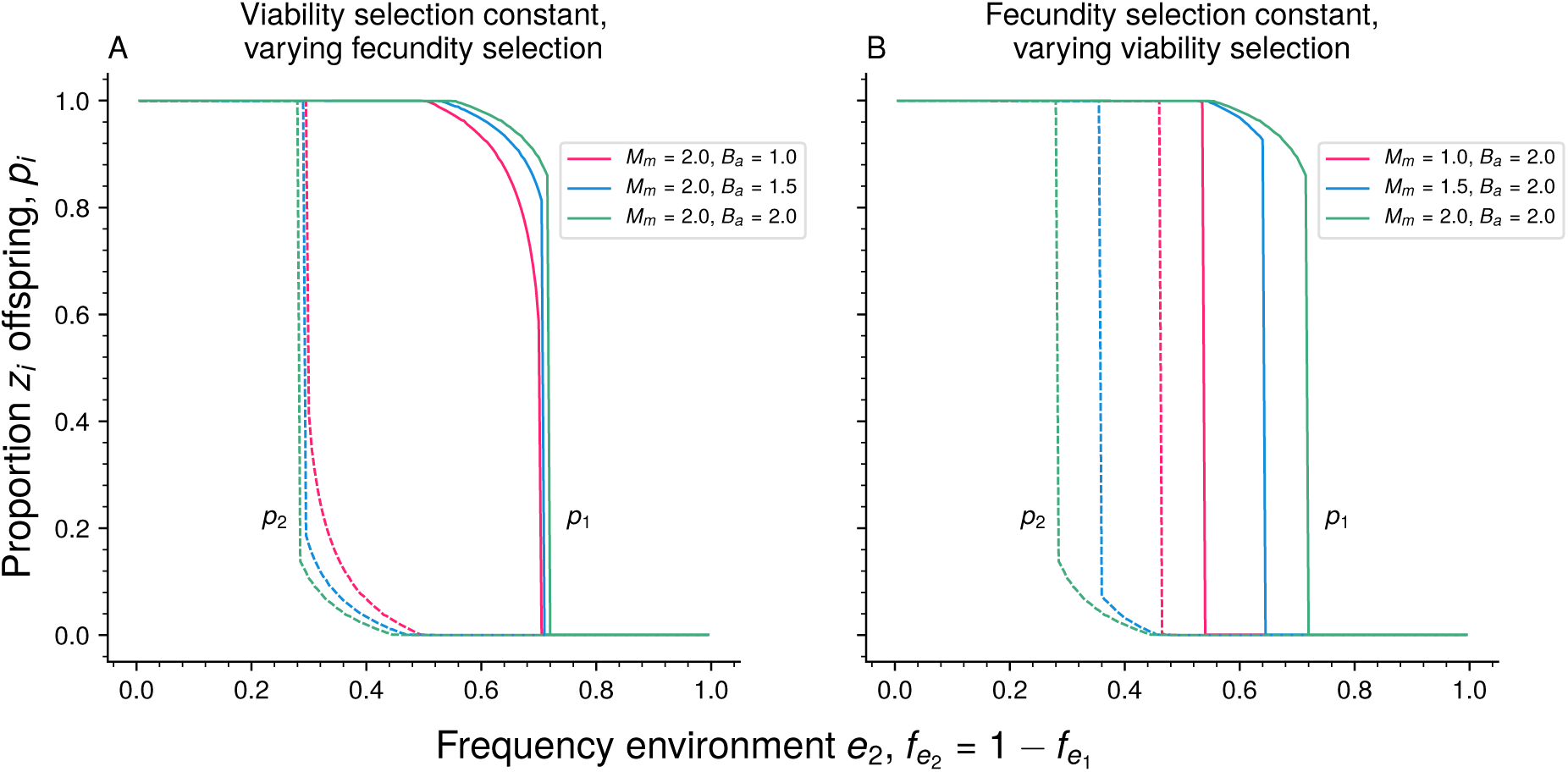
The evolution of parental effects (i.e., *p*_1_ ≠ *p*_2_: where dotted and solid lines do not overlap) in populations experiencing different combinations of viability selection and fecundity selection. Panel A: populations experience a constant, high level of viability selection *M*_m_ = 2 : *M*_a_ = 1 (viability selection on maladapted:adapted breeders) and varying levels of fecundity selection *B*_m_ = 1 : *B*_a_ varies (fecundity selection on maladapted:adapted breeders). When combined with viability selection, fecundity selection slightly enhances (rather than inhibits) the scope for parental effects. Panel B: populations experience a constant, high level of fecundity selection and varying levels of viability selection. The scope for parental effects increases with ever increasing levels of mortality selection. Parameters as in Figure 1.

## 4 Discussion

The current analysis shows that the mode of selection matters to the evolution of phenotypic inheritance: in spatially structured populations that experience fecundity selection, high-fidelity inheritance is favored with little scope for parental effects. By contrast, in populations which experience viability selection, the evolutionary scope for parental effects is substantially larger. Hence, life-history differences that affect the influence of fecundity versus viability selection may play an important role in the evolution of parental effects and phenotypic memory.

Why do we predict that viability and fecundity selection selection have these different consequences to the evolution of parental effects? Essential for the evolution of parental effects is that the parental phenotype evolves to be a reliable cue for offspring phenotype determination, which requires that the parental phenotype is correlated with the local environment (Kuijper & Hoyle, 2015; McNamara *et al.*, 2016; Dall *et al.*, 2015). Because viability selection causes locally adapted breeders to survive for longer than locally maladapted breeders, a breeder is more likely than not to have a phenotype matching the local environment, simply because it is being alive (McNamara & Dall, 2011). In other words, a correlation between phenotype and environment immediately builds up, because survival selection results in the enrichment of patches with phenotypes matching the local environment. This enrichment of locally adapted phenotypes is illustrated in Figure S5A: even when the distribution of offspring phenotypes is completely random (i.e., when *p*_1_ = *p*_2_ = 0.5), viability selection causes locally adapted phenotypes to prevail in each environment. Hence, in regimes of viability selection, parental phenotypes often are informative cues about their local environment, favoring the evolution of parental effects, as demonstrated by a large covariance between the parental phenotype and its environment (Figure S6A).

In regimes of fecundity selection, however, mortality rates of locally adapted and maladapted breeders are identical (because selection acts on the number of offspring produced rather than on survival). Hence, a breeder does not gain any additional information about its match to the local environment by simply being alive. A correlation between phenotypes and the local environment still develops, however, when two conditions are met: (i) more fecund, locally adapted parents produce most of the successfully established offspring in the local patch and (ii) the phenotypes of these successfully established offspring are correlated to the phenotypes of their parents. Although requirement (i) is a logical consequence of fecundity selection, note that correlations now only develop in the next generation (i.e., after successful offspring establishment), which is slower than in regimes of viability selection where correlations arise due to differential survival within one and the same generation. Hence, this not only means that it takes longer for correlations to build up again after the local environment has changed, but also that correlations are weakened by the establishment of any immigrant juveniles. Finally, the second requirement requires the presence of some inheritance fidelity, so that parental effects should already have evolved in the first place. Indeed, Figure S5B shows that when offspring have their phenotype randomly assigned, there is no parameter space where both phenotypes are associated with their respective environments simultaneously. By contrast, when there is some inheritance fidelity, a correlation develops, but it is still substantially weaker than in regimes of viability selection (see Figure S6B). Hence, in regimes of fecundity selection, parental phenotypes are less likely to be informative cues about their local environment, thus narrowing the scope for the evolution of parental effects.

The prediction that inheritance fidelity depends on the mode of selection has several interesting consequences for the current study of inheritance systems (e.g., Bonduriansky & Day, 2009; Jablonka & Raz, 2009; Jablonka & Lamb, 2010; Danchin *et al.*, 2011; Bonduriansky *et al.*, 2011). First of all, our study would predict that the scope for extended inheritance is generally weaker in longer-lived organisms, which are typically characterized by weak viability selection relative to fecundity selection during adult life (Stearns, 1992; Roff, 2002). For example, contemporary human populations exhibit little interindividual variation in adult survival, yet exhibit substantial variation in fecundity and fertility (Byars *et al.*, 2010; Stearns *et al.*, 2010; Courtiol *et al.*, 2012). Extrapolating from our model, we may therefore expect that selection in contemporary populations may largely disfavor inheritance mechanisms with relatively low fidelity, such as positive maternal effects or epigenetic inheritance mechanisms. Such lack of nongenetic effects may be indicative of a limited influence of the prenatal environment on future life expectancy, as found by a number of studies (Wells, 2007; Rickard & Lummaa, 2007; Hayward *et al.*, 2013; Wells, 2012a). However, we emphasize that while selection in contemporary populations could disfavor low-fidelity inheritance, evidence for nongenetic inheritance in humans may well be the result of past selection, during which viability selection was likely to be considerably higher (e.g., Wells, 2012b). In general, whether parental effects affect offspring viability or fecundity is thus important to take into account.

How to test the predicted relationship between parental effects and viability versus fecundity selection? One method of choice would be to experimentally evolve populations in fluctuating environments, where one contrasts lines that experience fluctuating selection on early fecundity with lines that experience fluctuating selection on longevity (e.g., through sampling offspring from individuals in later life). Experimental evolution in regimes that favor either fecundity or viability has been previously done in *Caenorhabditis* nematodes (e.g., Anderson *et al.*, 2011). Moreover, there is now accumulating evidence that parental effects can rapidly evolve in *Caenorhabditis* nematodes (Dey *et al.*, 2016; Lind *et al.*, 2019), making this an ideal model system to assess the evolution of parental effects in different life-history contexts. A key caveat is, however, that these experimental evolution studies have been performed on well-mixed populations, whereas our study suggests that a difference between viability and fecundity selection may only arise in spatially structured populations with limited dispersal and relatively small deme sizes. It would thus be interesting to consider the experimental evolution of transgenerational effects in such contexts, as has been done in *Caenorhabditis* within the context of local adaptation (Friedenberg, 2003; Teotónio *et al.*, 2017).

Our model highlights that differences in the type of selection acting on the population may be much more important to the evolution of parental effects than currently anticipated, yet there is substantial scope for future improvements. Foremost, our model focuses on the evolution of ‘cascading’ parental effects (McGlothlin & Galloway, 2013), where offspring phenotype determination depends on cues about the parental phenotype, rather than on more direct cues about the parental environment (environmental maternal effects (Lacey, 1998)), or cues about the parent’s genes (maternal genetic effects). In contrast to the current model, we would expect that the difference between fecundity and viability selection is unlikely to be important for environmental maternal effects. This is because offspring now receive a direct cue about the maternal environment, hence the development of a correlation between the maternal phenotype and the local environment through viability or fecundity selection becomes irrelevant. However, in the context of maternal genetic effects, we would expect differences between fecundity and viability selection to be similarly important as in the current model. This is because the maternal genotype only becomes informative to offspring when it correlates to the local environment (through differential survival of its bearers). As the current paper shows, viability selection has a much higher efficacy in developing such correlations. It would be welcome to expose the different types of parental effects to a systematic evolutionary analysis across different selective contexts: a welcome first step in this direction was undertaken recently by Proulx & Teotónio (2017) who compared maternal genetic and maternal environmental effects in the context of viability selection.

Another assumption of the current analysis is that parental effects affect binary phenotypes (*z*_1_ and *z*_2_), thus reflecting those traits which are dimorphic rather than continuous (Roff, 1996). It would be interesting to extend the current analysis to the evolution of parental effects on continuous traits, as done by quantitative genetics modeling when populations are well-mixed (e.g., Rivoire & Leibler, 2014; Kuijper & Hoyle, 2015) or are characterized by very high numbers of breeders per patch (Leimar & McNamara, 2015). However, given previous findings that binary and continuous models of parental effects often result in qualitatively similar outcomes (e.g., increased importance of maternal effects when dispersal is limited and when environments are autocorrelated), we would expect that the current conclusions are robust to contexts where traits are continuous rather than discrete.

Finally, for the sake of comparison with previous analyses on the evolution of parental effects (e.g., Proulx & Teotónio, 2017; McNamara *et al.*, 2016; English *et al.*, 2015), we have focused on a simple trait that is under stabilizing selection in a variable environment. Future models should consider, however, the evolution of these parental effects in the face of explicit life-history trade-offs, such as between offspring size and number (Smith & Fretwell, 1974; Krist, 2011), between reproductive effort and survival (Schaffer, 1974; Santos & Nakagawa, 2012) or both simultaneously (Winkler & Wallin, 1987). Existing theoretical predictions on offspring size versus number decisions have been highly valuable (reviewed in Rollinson & Rowe (2015)), yet these analyses only consider offspring phenotypes up and until juvenile survival (e.g., Smith & Fretwell 1974; Parker & Begon 1986; Kuijper & Johnstone 2013, but see Mangel *et al.* 1994). By contrast, a more rigorous analysis of parental effects would consider whether a maternal decision about sizes/numbers of her young subsequently affects those same decisions when made by her offspring and by later descendants (Pick *et al.*, 2019). Consequently, future analyses are needed that track the evolution of parental effects from life-history traits in parents to life-history traits in offspring. Particularly welcome would be a comparison between the evolution of parental effects which affect offspring reproductive effort versus survival in adulthood (viability selection) and parental effects that affect offspring size versus number decisions (fecundity selection).

## Acknowledgments

This study was funded by a Leverhulme Trust Early Career Research Fellowship (ECF 2015-273) awarded to BK. This work has made use of the Carson computing cluster at the University of Exeter’s Environment and Sustainability institute.

