## Supplementary Figures for "The evolution of parental effects when selection acts on fecundity versus viability"

*Online Supplement*

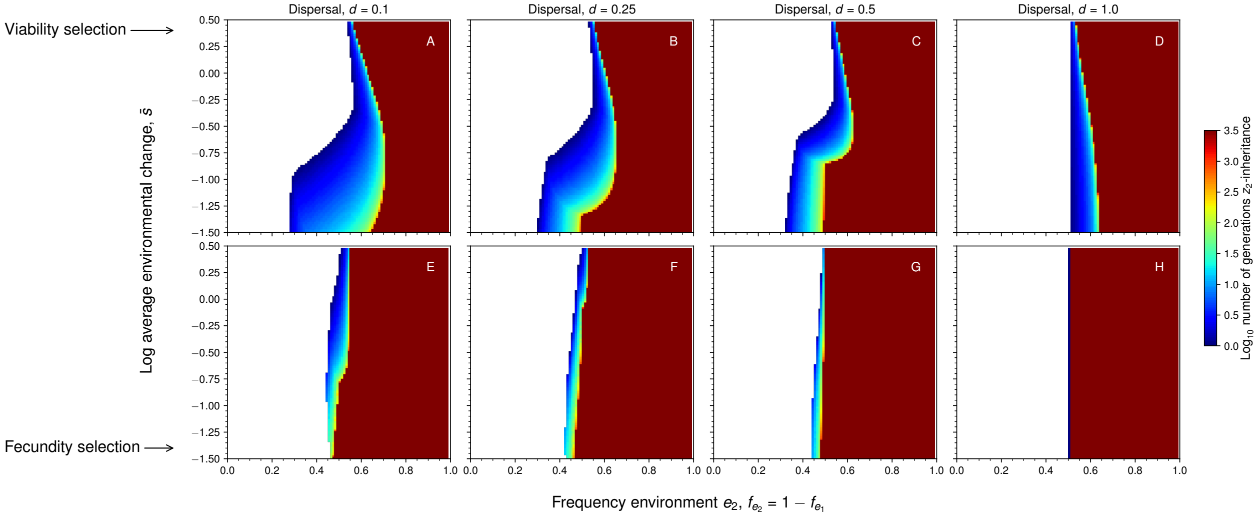


**Figure S1**

**Figure S1** Fecundity selection (Panels E-H), but not viability selection (Panels A-D) substantially reduces the parameter space where parental effects lead to phenotypic inheritance of phenotype $z_2$. See Figure 2 in the main text for phenotype $z_1$.

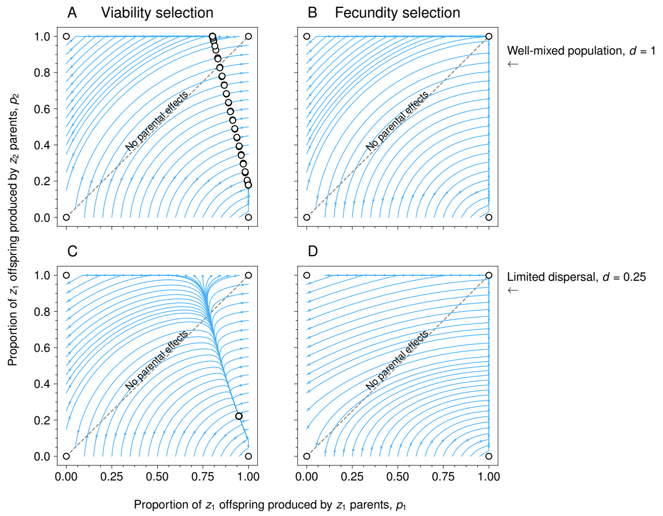


**Figure S2**

**Figure S2** Evolutionary trajectories of phenotype determination strategies $p_1$ and $p_2$ for viability selection (panels A, C) and fecundity selection (panels B, C). The first row of the figure depicts a scenario where all juveniles disperse ($d = 1$) and the second row populations with limited dispersal ($d = 0.25$). We consider an environment that changes at modest rate $\bar{s} = -1$ and where $e_1$ patches are more common than $e_2$ patches ($f_{e_1} = 1 - f_{e_2} = 0.6$).

**Fecundity selection, panels B, D:** For the environment considered in the current figure, fecundity selection always result in outcomes where parental effects are absent ($p_1 = p_2 = 1$; panels A, C), resulting in a $z_1$ monomorphism. See also Figure 2 in the main text.

**Viaibility selection, panel A:** In case of viability selection, we find that $p_1$ and $p_2$ evolve to a line of equilibra, but only when $d = 1$. The line of equilibria implies that parental effects are very likely to be present (as the point $p_1 = p_2$ where parental effects are absent is only one of infinitely many points along the line), despite the fact that there is no correlation between the parental phenotype and the offspring's environment (as offspring migrate to a random patch). Moreover, once a population has arrived at the line of equilibria, genetic drift along the line of equilibria can lead to between-population differentiation in the values of parental effects. Why do we find this line of equilibria when $d = 1$? Due to the global distribution of $e_1$ (frequency 0.6) and $e_2$ patches (frequency 0.4), an overall mixture of $z_1$ and $z_2$ offspring (biased towards $z_1$ offspring because $e_1$ is more common) is selectively favored. However, such a global mixture can be produced by a multitude of combinations of $p_1$ and $p_2$, thus resulting in a line of equilibria. By contrast, when dispersal is limited, there is no global mixture of $z_1$ and $z_2$ offspring that is favored in regimes of mortality selection, as local patches in environmental state $e_1$ favor a different mixture of offspring phenotypes relative to local patches in environmental state $e_2$. Hence, the line of equilibria collapses into a single point (see Figure S2C).

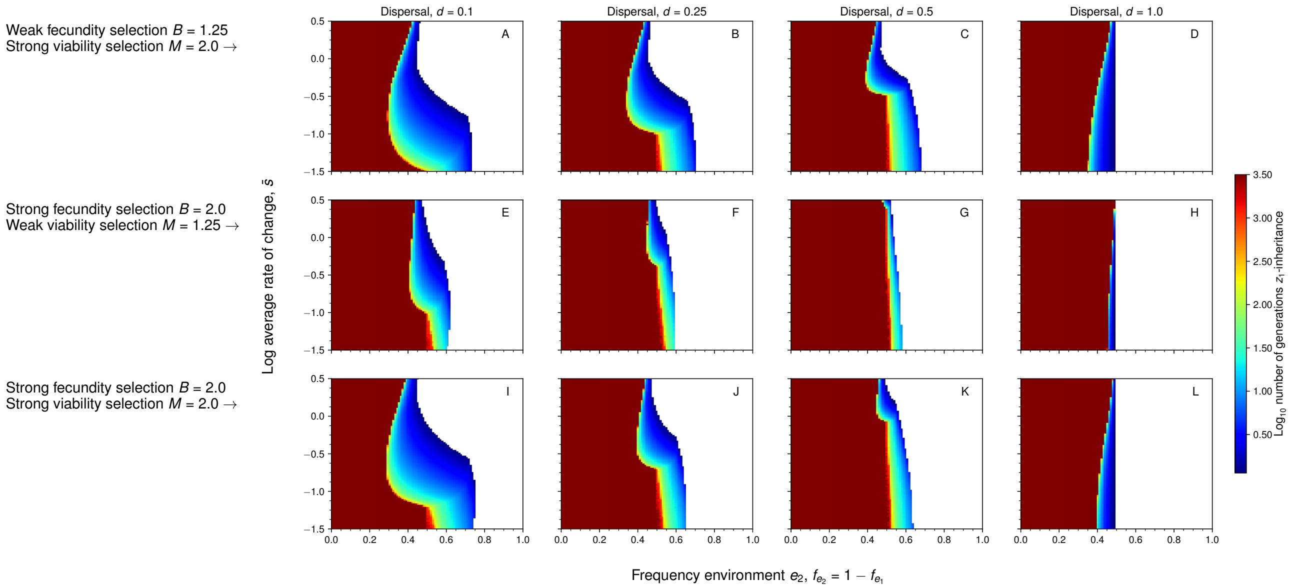


**Figure S3**

**Figure S3** Combinations of fecundity selection and viability selection affecting the duration of inheritance. When viability and fecundity selection act simultaneously, we find that the evolution of parental effects (and hence the duration of inheritance) is largely shaped by viability selection and does not seem to be reduced by fecundity selection. Rather, when viability selection is already present, fecundity selection can slightly enhance the parameter space in which intermediate levels of inheritance occur (e.g., panels I - L where both viability and fecundity selection are strong). Parameters are identical to Figure 2 in the main text.

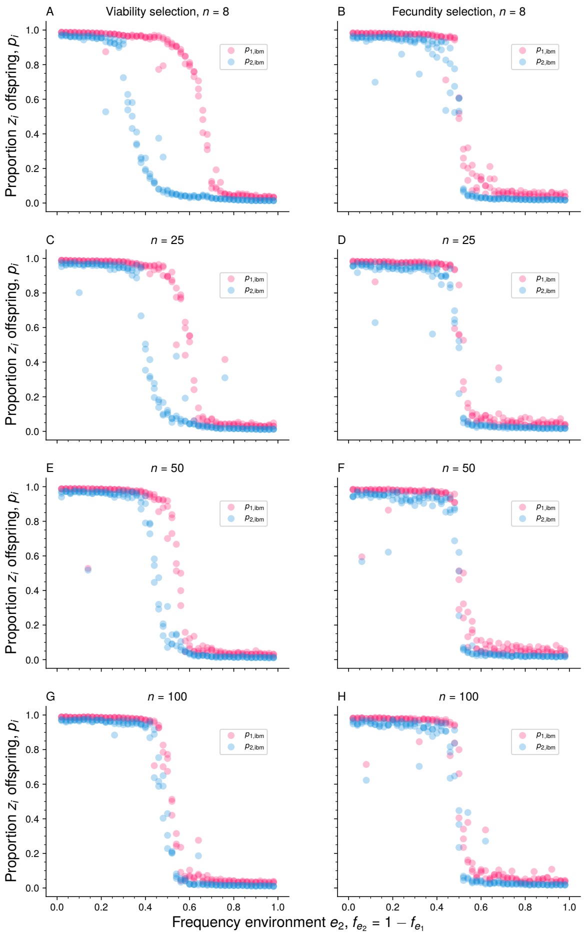


**Figure S4**

**Figure S4** Individual-based simulations when varying the number of breeders $n$ per patch. Overall, we find that parental effects are more likely to evolve in populations experiencing viability but not fecundity selection, unless patch sizes become vary large (e.g., $n = 50, 100$) in which case no parental effects evolve. Parameters: $d = 0.25$.

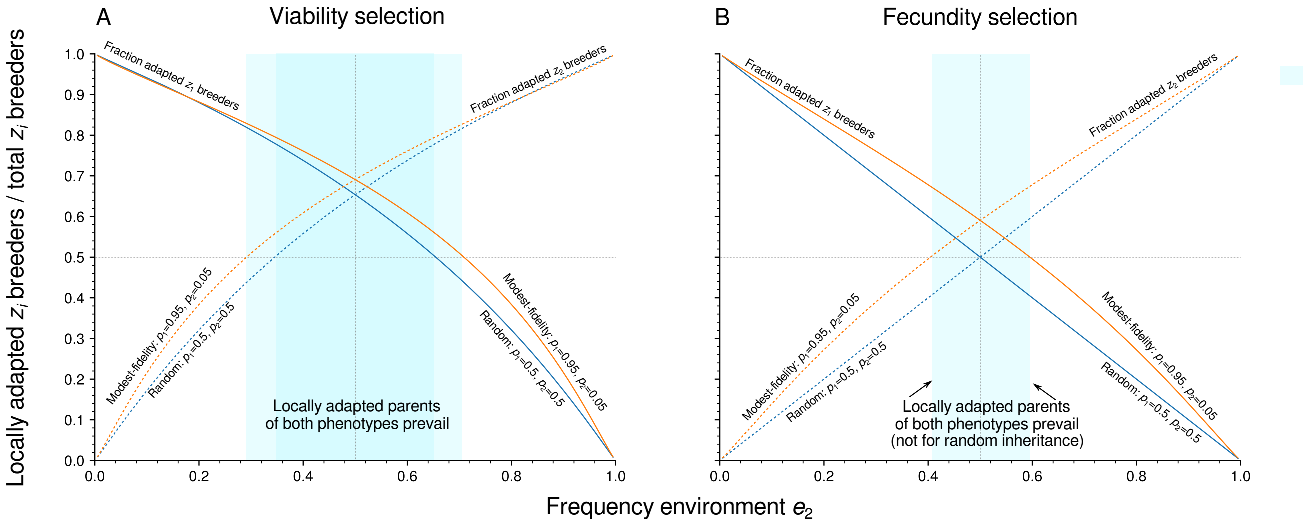


**Figure S5**

**Figure** S5 Effect of viability and fecundity selection on the fraction of individuals of each phenotype $z_i$ (relative to the total number of $z_i$ individuals) who are locally adapted.

Panel A (viability selection): even in the face of random phenotype determination (blue solid and blue dotted lines; $p_1 = p_2 = 0.5$), viability selection still causes patches of both environments $e_1$ and $e_2$ to become enriched with $z_1$ and $z_2$ individuals respectively, as the fraction of locally adapted $z_1$ and $z_2$ individuals exceeds 0.5 for a broad range of the parameter space (cyan-colored regions). This association between both phenotypes and their environments is even larger when parents can pass on their locally adapted phenotype to their offspring (orange lines).

Panel B (fecundity selection): when phenotype determination is random (blue and solid dotted lines), fecundity selection does not cause $z_1$ and $z_2$ to become any more enriched in their local environments relative to the probability that a $z_i$ finds itself on a $e_i$ patch by chance (i.e., the frequency of a $e_i$ environment). Only when locally adapted (and hence more fecund) parents leave more locally adapted descendants to the local patch, will there be enrichment of environments with locally adapted individuals. However, this enrichment occurs only in roughly a third of the parameter space relative to that of viability selection. Parameters as in Figure 1 in the main text.

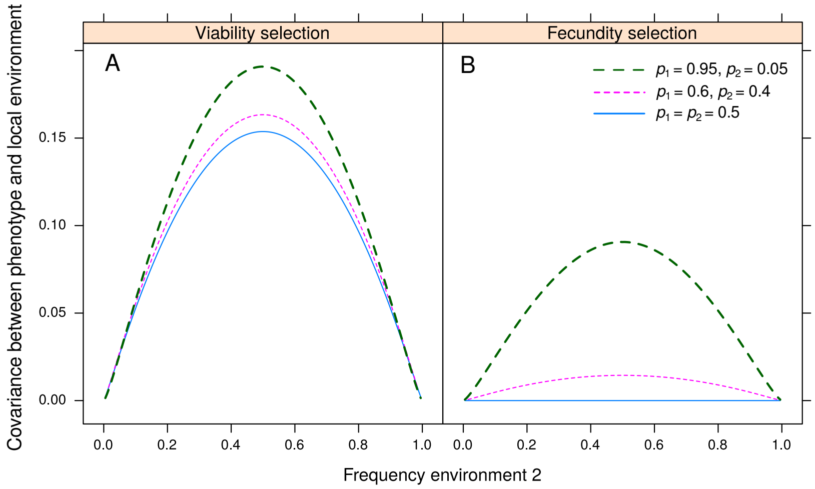


**Figure S6**

**Figure S6** Effect of viability (panel A) and fecundity selection (panel B) on the covariance between phenotypes and their local environments for different fixed values of the phenotype determination strategies $p_1$ and $p_2$. Overall, covariances between phenotype and environment are always weaker for fecundity selection and are 0 when $p_1 = p_2 = 0.5$ (random phenotype determination). By contrast, covariances are always nonzero, even for random phenotype determination in populations experiencing viability selection. Coviarances between phenotype and environment $\text{cov}(e_i, z_i) = \text{E}[e_i z_i] - \text{E}[e_i]\text{E}[z_i]$ are calculated by giving individuals of phenotype $z_1$ and $z_2$ the indices 1 and 0 and similarly by giving environments $e_1$ and $e_2$ the indices 1 and 0. We then have $\text{E}[e_i z_i] = \sum_{n_{z_1}=0}^{n} f_{n_{z_1} e_1} n_{z_1}$, $\text{E}[e_i] = \sum f_{n_{z_1} e_1}$ and $\text{E}[z_i] = \sum_{n_{z_1}=0}^{n} \sum_{e_i \in \{e_1, e_2\}} f_{n_{z_1} e_i} n_{z_1}$. Parameters as in Figure 1 in the main text.
